# From predictive models to cognitive models: Separable behavioral processes underlying reward learning in the rat

**DOI:** 10.1101/461129

**Authors:** Kevin J. Miller, Matthew M. Botvinick, Carlos D. Brody

## Abstract

Cognitive models are a fundamental tool in computational neuroscience, embodying in software precise hypotheses about the algorithms by which the brain gives rise to behavior. The development of such models is often a hypothesis-first process, drawing on inspiration from the literature and the creativity of the individual researcher to construct a model, and afterwards testing the model against experimental data. Here, we adopt a complementary approach, in which richly characterizing and summarizing the patterns present in a dataset reveals an appropriate cognitive model, without recourse to an *a priori* hypothesis. We apply this approach to a large behavioral dataset from rats performing a dynamic reward learning task. The revealed model suggests that behavior in this task can be understood as a mixture of three components with different timescales: a quick-learning reward-seeking component, a slower-learning perseverative component, and a very slow “gambler’s fallacy” component.

## Introduction

A fundamental goal of cognitive neuroscience is to understand the algorithms by which the brain gives rise to behavior. One of the basic tools used in this pursuit is cognitive modeling. This involves constructing software agents which are capable of performing tasks, and tuning them such that their behavior matches as closely as possible the behavior of human or animal subjects. It also involves treating the algorithms implemented by these agents as precise hypotheses about the neural algorithms implemented by the brain to perform those same tasks (Corrado & Doya, 2007; Daw, 2011; O’Doherty et al., 2007; Wilson & Collins, 2019). Cognitive models have been used to shed light on the neural mechanisms of many aspects of cognition, including learning (Daw & Doya, 2006; Lee et al., 2012), memory (Norman et al., 2008), attention (Heinke & Humphreys, 2005), and decision-making (Gold & Shadlen, 2007; Hanks & Summerfield, 2017; Sugrue et al., 2005).

Evaluating a cognitive model requires considering two fundamentally different criteria. The first is whether it is accurately able to predict behavioral data. This predictive accuracy can be quantified, and a number of tools exist for comparing it objectively among models (e.g. cross-validation; for overview see Gelman et al., 2013, Chapter 7). The second is whether it is plausible that the brain implements similar algorithms to the model. Though aspects of this can be quantified – many researchers agree that simpler models are preferable, and tools exist for quantifying model simplicity (but see Gelman & Rubin, 1995; Raftery, 1995; Wagenmakers et al., 2010) – it is ultimately a subjective evaluation that relies on outside knowledge about the psychology and biology of the system.

The development of new cognitive models often focuses first on cognitive plausibility. Researchers instantiate plausible hypotheses as software agents, test them against experimental data, and refine them when necessary to produce a better fit. One popular and successful approach, for example, draws inspiration from optimality, looking to algorithms which are provably the best solutions that exist for a particular problem (Gold & Shadlen, 2002; Griffiths et al., 2015; Körding, 2007). Another draws inspiration from artificial intelligence, looking to algorithms which are successful in practice at solving a wide variety of computational problems (Daw & Doya, 2006; Dayan & Niv, 2008). These and similar approaches have been extremely productive, but are subject to two important limitations. The first is that they typically consider a relatively small number of models. They are typically able to determine that a particular model is the best of a set, but not that it is good in an absolute sense (Daw, 2011). The second is that the development of new models depends on inspiration from adjacent fields or on the creativity of the individual researcher, limiting the pace of hypothesis generation.

Here, we adopt an alternative approach, which focuses first on predictive accuracy. This approach begins with a model that, given a particular characterization of the behavioral task, is maximally flexible. This model is not itself plausible as a cognitive hypothesis, but provides a first characterization of the patterns present in the dataset, as well as a standard against which to measure the predictive accuracy of other models. The approach continues with a process of successive model reduction: patterns are identified in the fit parameters of a more-flexible model, and a less-flexible model is proposed that embeds those patterns into its structural assumptions. If the predictive accuracy of the less-flexible model is similar to that of the more-flexible one, it is itself adopted as a target for further reduction.

This approach requires fitting highly flexible models, with large numbers of free parameters, and this requires in turn the collection of large datasets. In the present paper, we took advantage of high-throughput methods for studying decision-making in rats (Brunton et al., 2013; Erlich et al., 2011) to collect a large behavioral dataset from a classic reward-guided learning task, the “two-armed bandit” task (Daw et al., 2006; Ito & Doya, 2009; Kim et al., 2009; Samejima et al., 2005). Our approach of successive model reduction resulted ultimately in a relatively compact model, with just seven free parameters, that captures the major patterns present in rat behavior on this task. This model can be viewed as a cognitive model, describing a candidate for the neural algorithm implemented by the rat brain to solve the task. When viewed in this way, it suggests that behavior is the product of three distinct mechanisms – a reward-seeking mechanism that considers only the most recent few trials, and tends to repeat choices that led to rewards and switch away from those that did not; a perseverative mechanism that considers a slightly longer history, and tends to repeat choices regardless of their outcomes; and a “lose-stay” mechanism that considers a much longer history, and tends to repeat choices that did not lead to rewards. Each of these components was present consistently across rats, with rat-by-rat variability in their precise strengths and time constants. The first two components (reward-seeking and perseverative) are broadly consistent with existing theory, and represent refinements to existing models; the last component (lose-stay) is to our knowledge unexpected.

## Results

### Task

Rats performed a probabilistic reward learning task (the "two-armed bandit" task; Ito & Doya, 2009; Kim et al., 2009; Sul et al., 2010) in daily sessions (20 rats; 1,946 total sessions; 1,087,140 total trials). In each trial, the rat first entered a central noseport, then selected between a noseport on the left and one on the right (“choice ports”), and finally received either a water reward or a reward omission (Figure 1a). The probability of reward at each choice port evolved slowly over time according to a random walk (Figure 1b). Task performance required ongoing learning about the current reward contingencies, and selecting the port with the higher reward probability. The rats performed well at this task, tending to select the nose port that was associated with a higher probability of reward (Figure 1c). Importantly, rats’ performance did not match that of an agent incorporating a Bayesian ideal observer (Figure 1c, see Methods), meaning that the patterns in their behavior are not purely a function of the task itself, but rather of the strategy with which the rats approach it.

**Figure One:**
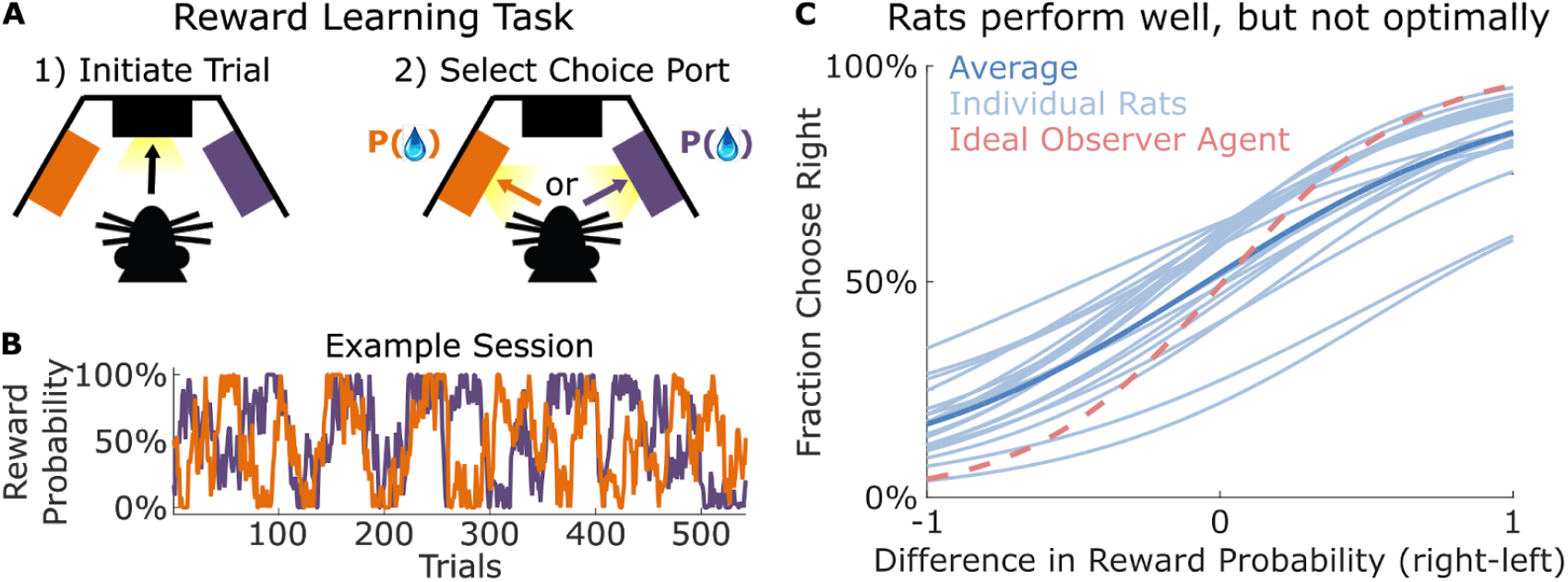
Two-armed bandit task for rats. **A)** On each trial, rat selects one of two possible choice ports. **B)** Probability of reward at each port evolves over time according to an independent random walk. **C)** Rats tend to select the port with the higher reward probability. Their performance does not match that of an ideal observer agent.

### Unconstrained Models

First, we consider predictive models which characterize in a very general way the relationship between a rat’s choices and the recent history of choices and rewards. For these models, and those that follow, we choose to model the rats choice on each trial, using as predictors the recent history of choices and their outcomes. These “unconstrained” models can be thought of as look-up tables, where each entry gives the probability that the rat will choose left vs. right following a particular history of *N* past choices and rewards (n-markov models; Ito & Doya, 2009). Specifically, we label the choice made (left or right) and the outcome received (reward or omission) on trial *t* as *C_t_* and *O_p_* respectively, and we define the history, *H,* as matching between a pair of trial if the choices and outcomes that preceded those trials are identical:

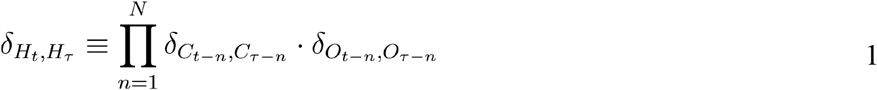

where *δ* is the Kroenecker delta function, which takes on a value of one when its arguments match and zero otherwise, and *N* is a hyperparameter governing how many past trials are considered by the model. The *δ_H_t_,_H_τ___* variable takes on a value of one only when the *N* choices and outcomes preceding trial *τ* match exactly with those preceding trial *τ.* The model’s predicted choice probability for each port is equal to the fraction of trials with matching histories in which the rat selected that port:

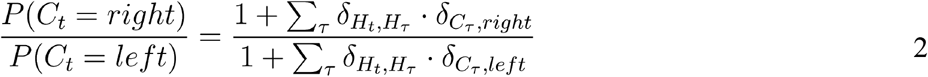

Predicted choice probabilities for *N*=1 and *N*=2 are shown in figure two for an example rat, and in figure S1 for all rats. The unconstrained model introduces no assumptions about the way in which recent past choices and outcomes influence future behavior. The cost of this flexibility is a very large number of free parameters: one per possible history *H*. Each trial within that history can be one of four types (left/right choice, reward/omission outcome), yielding 4^*N*^ parameters per rat. Fora given dataset, it will be possible to obtain useful estimates of these parameters only up to some value of *N*. For larger values, there will histories that repeat only rarely, resulting in a small number of nonzero entries in *δ_H_t_,_H_τ___* and a poor estimate of choice probability. A result of this is overfitting, in which a model fit to a particular dataset provides a better quality of fit to that dataset than to a separate dataset generated in the same way. We evaluate the performance of the models by computing a normalized likelihood score (Daw, 2011), using two-fold cross-validation. Specifically, we estimate one set of parameters using only the even-numbered sessions from a particular rat and another using only the odd-numbered sessions, then evaluate each set of parameters both on the sessions used to estimate it (“training dataset”) and on the other sessions (“test dataset”). We find that test-set likelihood is similar to training-set likelihood for models with short history windows (*N* ≲ 4), indicating relatively little overfitting (example rat: figure 2C, all rats: figure S1). For models with longer history windows (*N* ≳ 4), testing-set and training-set likelihood diverge, indicating that substantial overfitting has occurred. The training-set likelihood provides a lower bound on the true quality of model fit, while the testing-set likelihood provides an upper bound. In the region where these are similar, we have a reliable estimate of model quality, which we can use to compare against other models. This estimate provides a ceiling on the performance that can be achieved by any model that considers only recent choices and rewards, since the unconstrained model is the most flexible such model.

**Figure Two:**
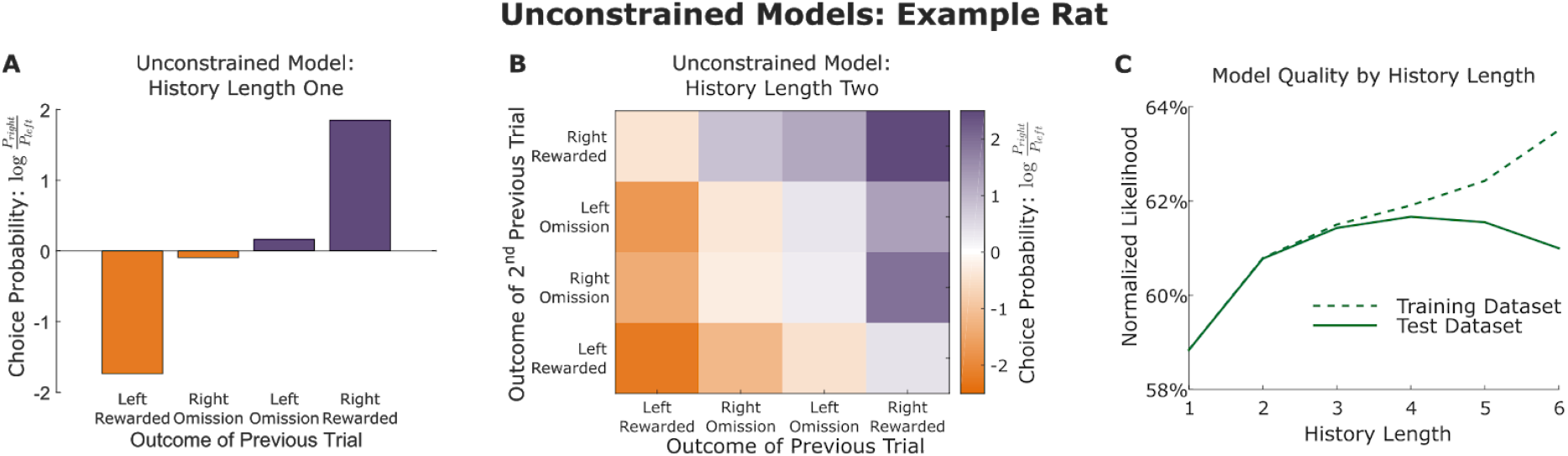
Unconstrained models. **A)** Unconstrained model considering one previous trial, fit to a dataset from an example rat. This model comprises four parameters, each giving the log choice probability ratio immediately following each of the four possible trial types. **B)** Unconstrained model considering two previous trials, fit to the same example rat. This model comprises sixteen parameters, each giving the log choice probability ratio immediately following a particular pair of trial types. **C)** Quality of fit for unconstrained models considering different history lengths. Normalized likelihood was computed using cross-validation, and likelihoods for both the fit datasets (training dataset) and the held-out datasets (testing dataset) are plotted. Likelihoods are similar for both datasets when the order is small.

### Linear models

Next, we consider predictive models which mitigate overfitting by introducing an assumption about the way in which past trials choices and outcomes affect future behavior. In particular, these models assume that each past trial exerts an influence on future choice that is independent of the influence of all other past trials. The influence of each past trial is summed, and this sum is used to compute choice probability. The most general models in this family fit the relationship of the sum to choice probability (linear-nonlinear-poisson models; Corrado et al., 2005), but we consider here models which additionally assume that this relationship is well-approximated by a logit function (logistic regression models; Lau & Glimcher, 2005). For our task, the most general of these can be written as:

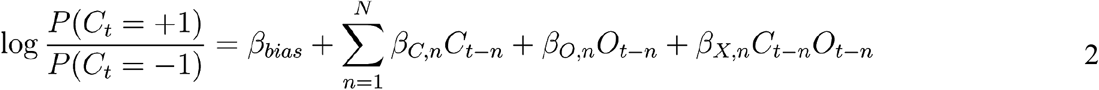

where *C_t_* is the choice made on trial *t* (coded as +1 for right and −1 for left), *O_t_* is the outcome received (+1 for a reward and −1 for an omission), and the *βs* are fit weighting parameters. Like the unconstrained models, these models consider a history of *N* recent trials, but unlike them, they do not assign a parameter to each possible history. Instead, they assign a set of weights to each slot within that history, quantifying the effect on upcoming choice when a trial of each type occupies that slot. In our model, these weights are organized into three vectors: *β_X_*, which quantifies a “reward seeking” pattern in which the animal tends to repeat choices that have led to rewards and to switch away from choices that have led to omissions; *β_C_*, which quantifies a “choice perseveration” pattern in which the animal tends to repeat past choices without regard to their outcomes; and *β_O_*, which would quantify any pattern in which outcomes affect choice without regard to the side on which they were delivered (the symmetry of the task suggests this term is likely to be near-zero). The model also contains a bias parameter *β_bias_*, which quantifies any fixed left/right choice bias.

Fits of the linear model reveal large and consistent effects of reward seeking and choice perseveration, and relatively weak and inconsistent main effects of outcome (example rat: Figure 3A; all rats: Figure S2). These fits were relatively resistant to overfitting: normalized likelihoods for the testing and training datasets diverged from one another at much larger *N* than those of the unconstrained model (Figure 3B, compare to Figure 2B). This resistance to overfitting comes at the cost of the assumption that past outcomes contribute independently to current choice. One way to test this assumption is to compare the cross-validated likelihood scores of the unconstrained and the linear models directly: if the additional flexibility of the unconstrained model allows it to achieve a higher score, then meaningful nonlinear interactions must exist in the dataset. We find that the linear and the unconstrained model earn similar likelihood scores up to *N* ≈ 4, and that for larger *N* the linear model achieves larger scores, as the unconstrained models begin to overfit (Figure 3C). At no value of *N* do the unconstrained models outperform the linear models. This partially validates the assumptions of the linear models. It rules out strategies incorporating nonlinear interactions within the most recent few trials (e.g. “repeat the choice until getting two omissions in a row, then switch”), but not those incorporating longer-term nonlinear interactions (e.g. “if the most recent choice matches the choice from six trials ago, repeat it”). These results also provide a standard against which to evaluate other models which incorporate both its assumptions as well as others. Such models will be less flexible than the linear models, and the performance of the linear models provides a ceiling on the predictive accuracy that can be expected from them.

**Figure Three:**
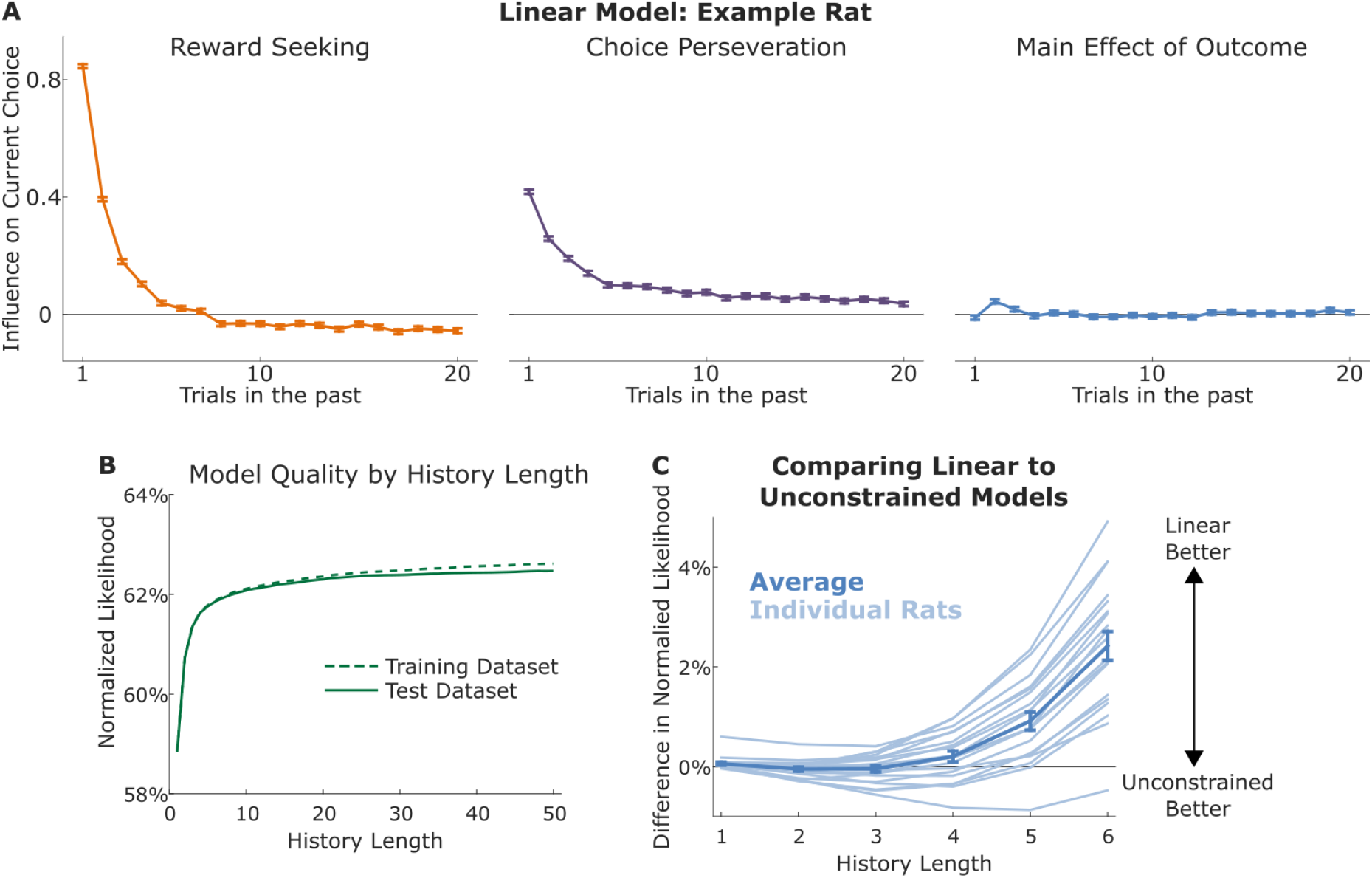
Linear Models. **A)** Linear model considering a history of length twenty, fit to the example rat. The model consists of three sets of weights characterizing the influence of past trials on current choice. Reward-seeking weights (left) characterize the tendency to repeat choices that led to rewards and switch away from choices that led to omissions; perseveration weights (middle) characterize the tendency to repeat past choices regardless of their outcomes; outcome weights (right) characterize the tendency for outcomes (reward/omission) to influence choice regardless of which choice preceded them. **B)** Quality of model fit for the example rat, as a function of the number of past trials considered. Normalized likelihood for both the fit datasets (“training dataset”) and the held-out datasets (“testing dataset”) are plotted. Likelihoods are similar for both datasets, even when considering large numbers of past trials (compare to Figure 2C). **C)** Difference in normalized cross-validated likelihood between linear and unconstrained models considering the same history length. Quality of fit of the linear model approximates or exceeds that of the unconstrained model for all history lengths.

### Mixture-of-Exponentials Models

Inspecting the fit weights of the linear models (Figure 3a, Figure S2), several patterns are apparent. For all rats, reward-seeking weights are large and positive for the most recent few trials (Figure 3A, left), while perseverative weights were smaller, but extended further back in time (Figure 3A, middle). Outcome weights were small and showed patterns that were inconsistent across rats (Figure 3A, right). We sought to capture these patterns using “mixture of exponentials” models, each comprising a mixture of exponential components.

Each exponential component exhibits either reward-seeking or perseveration, and was characterized by an update rate *α*, determining the component’s timescale, as well as by a weighting parameter *β*, determining the strength of its influence on behavior. These parameters govern the evolution of a hidden variable, which we denote *V* for reward-seeking components and *H* for perseverative components. These hidden variables were set to zero on the first trial of each session and updated after each trial according to the following rules:

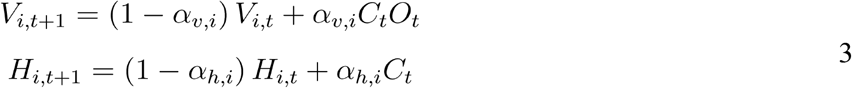

where *C_t_* and *O_t_* are the choice made and outcome received on trial *t* (coded +1 or −1, as above). Choice probability on each trial was determined by summing the influence of each of the exponential processes, as well as overall bias:

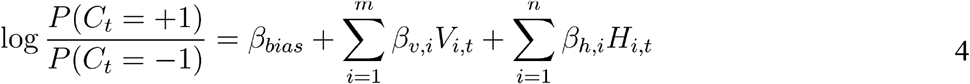

where *m* and *n* are hyperparameters controlling the number of reward-seeking and of perseverative components, and *β_v_* and *β_h_* are vectors of weighting parameters (of length *m* and *n* respectively).

We computed the quality of model fit as a function of the number of components included (Figure 4A). We found that the dataset was best explained by a mixture of exactly two reward-seeking and two perseverative components. Removing a component of either type resulted in a lower quality of fit, whether the removed component was reward-seeking (mean decrease in normalized likelihood of 0.21 percentage points, standard error 0.04, p=2×10^-4^, signrank test) or perseverative (mean 0.31, sem 0.05, p=1×10^-4^). Allowing the model to use an additional component of either type did not meaningfully improve quality of fit, whether the additional component was reward-seeking (mean improvement 0.01 percentage points, sem 0.006, p=0.14) or perseverative (mean improvement 0.01 percentage points, sem 0.02, p=0.07). We also found that a model with two reward-seeking and two perseverative components narrowly outperformed the best linear model for each rat (mean difference in normalized likelihood 0.16 percentage points, sem 0.04, p=0.001; Figure 4A).

**Figure 4:**
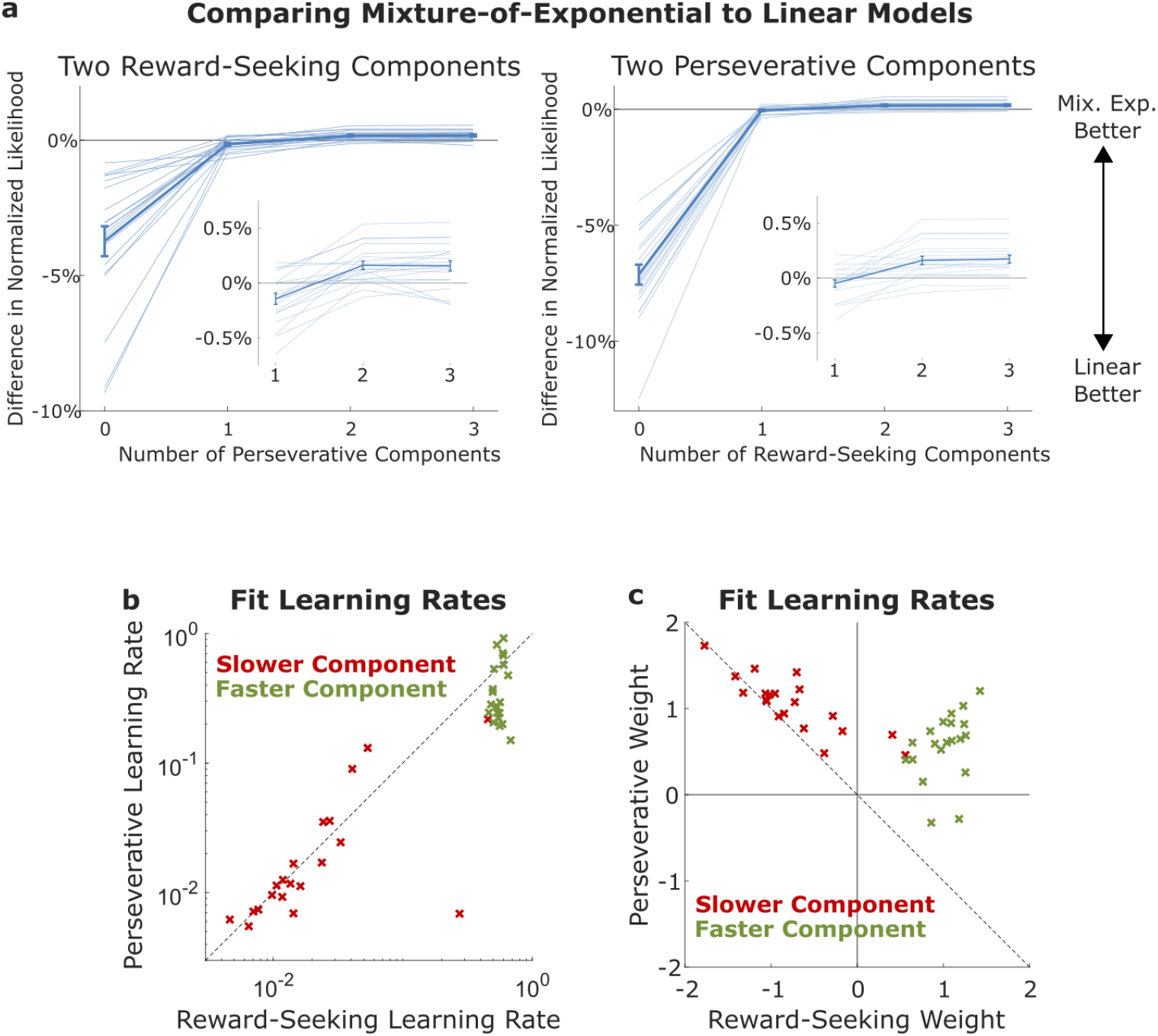
Mixture-of-Exponentials Models. **A)** Difference in normalized cross-validated likelihood between mixture-of-exponentials models with different numbers of exponential components and the best linear model for each rat. Left: Model with two reward-seeking components and different numbers of perseverative components. Left inset: Magnified view showing normalized likelihood differences for models with one, two, or three reward-seeking components. Right: Models with two perseverative components and different numbers of reward-seeking components. Right inset: Magnified view showing normalized likelihood differences for models with one, two, or three reward-seeking components. Quality of fit increases until two processes of each type are included, and then does not continue to increase.**B)** Learning rates of a mixture-of-exponentials model with two processes of each type fit to each rat. Learning rates of the slower perseverative and the slower reward-seeking component are strongly correlated, with a slope close to one. **C)** Weights of a mixture-of-exponentials model with two processes of each type fit to each rat. Weights of the slower perseverative and the slower reward-seeking component are strongly correlated, with a slope close to negative one.

Having established that it provides a good match to our rats’ data, we fit a model containing two reward-seeking and two perseverative components to the full dataset for each rat, and examined its fit parameters. We label each component “faster” or “slower” according to whether it had the larger or the smaller learning rate of the two components of its type. To reveal patterns among the fit parameters, we plot the learning rates (Fig 4B) and weights (Fig 4C) of the slower reward-seeking component against those of the slower perseverative component (red), and similarly the parameters of the faster reward-seeking component against those of the faster reward-seeking component (green). The learning rate of the slow reward-seeking component and of the slow perseverative component were highly correlated (r^2^=0.50, p<1×10^-3^), and many of them were nearly equal (Fig 4B, red points close to diagonal dotted line). This suggests a further model reduction: these two slow components may reflect the same underlying process. Similarly, the fit weight of the slow reward-seeking component and of the slow perseverative component were highly correlated (r^2^=0.68, p<1×10^-5^), and many of them were nearly equal in sign, though opposite in magnitude (Fig 4C, red points close to diagonal dotted line). This suggests that the slow underlying process may exhibit equal-and-opposite influences of reward-seeking and perseveration. The parameters associated with the two fast components showed little correlation (r^2^=0.04, p=0.68 for learning rates; r^2^=0.13, p=0.12 for weights), suggesting that each reflects a separate underlying process (Figure 4B and 4C, green points show no clear structure).

In sum, fits of the mixture-of-exponentials models indicate that rat behavior can be well-captured by a model containing two reward-seeking components and two perseverative components. Fit parameters suggest that the slower reward-seeking and slower perseverative components might reflect a common underlying process containing equal-and-opposite influences of each.

### Cognitive Model

Finally, we sought to capture the behavioral patterns revealed by the mixture-of-exponentials models using a simpler model, and to express this model using notation typical of cognitive models. We write this as a mixture-of-agents model, in which each rat is modeled as a mixture of three independent agents following different strategies: a reward-seeking agent, a perseverative agent, and an agent influenced by each pattern in an equal-but-opposite way.

The reward-seeking agent represents a single internal variable *V.* This quantity is initialized to zero at the beginning of each session, and updated after each trial according to the following rule:

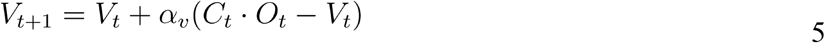

This is equivalent to an exponential reward-seeking process, as in equation 3. The variable *V* is updated towards a value of+1 following rewarded left choices (*C_t_*=1; *O_t_*=1) and unrewarded right choices (*C_t_* =−1; *O_t_* =−1), and updated towards a value of −1 following rewarded right choices (*C*_t_ =-1; *O_t_* =1) and unrewarded left choices (*C_t_*= 1; *O_t_*=−1). It represents the reward-seeking agent’s current relative preference for the right vs. the left choice port. This mechanism is similar to error-correcting mechanisms found in many animal learning and reinforcement learning theories (Bush & Mosteller, 1953; Rescorla & Wagner, 1972; Sutton & Barto, 2017), though different in that these models typically track the expected values of different actions independently, whereas the reward-seeking agent tracks only a single variable, which is affected by outcomes at both ports.

The perseverative agent represents a single internal variable *H,* which is initialized to zero at the beginning of each session, and updated after each trial according to:

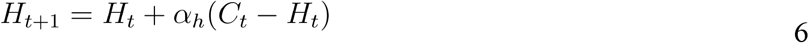

This is equivalent to an exponential perseverative process, as in equation 3. The variable *H* is updated towards +1 following left choices, regardless of reward, and towards a value of −1 following right choices. It represents a recency-weighted estimate of how often each port has been chosen. This can be thought of as a process of Hebbian plasticity that strengthens the representations of recently taken actions – a mechanism which has been proposed to underlie the formation of habits (Ashby et al., 2010; Miller et al., 2019).

The final agent represents an internal variable *G.* It is initialized to zero at the beginning of each session, and updated after each trial according to:

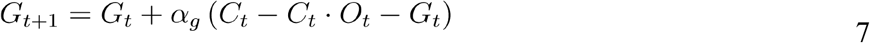

This is equivalent to an exponential process containing positive perseveration balanced by equal negative reward-seeking. Following rewarded trials (*O_t_*= 1), these influences cancel one another, and *G* is updated towards 0. Following unrewarded trials (*O_t_*= −1), these influences add to one another, and *G* is updated towards either +2 (left choices, *C_t_*= 1)or −2 (right choices, *C_t_*= −1). These updates result in *G* tracking a recency-weighted average of the relative number of losses incurred at each port. This pattern is reminiscent of the “gambler’s fallacy”, in which human subjects seem to believe that a long sequence of losing outcomes predicts that a winning outcome is likely to follow (Oskarsson et al., 2009; Tversky & Kahneman, 1971).The choice made by the model on each trial is determined by a weighted average of *V*, *H*, and *G*, as well as bias:

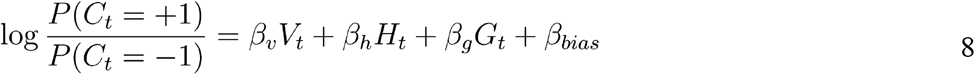

The complete cognitive model has seven parameters: three learning rates (*α_v_*, *α_h_* and *α_g_*) and four weights (*β_v_*, *β_h_*, *β_g_*, and *β_bias_*). It can be viewed as a simplification of the four-component mixture-of-exponentials model, with parameter equivalencies given by Table 1. Like this model, the cognitive model also narrowly outperforms the best linear model in terms of cross-validated likelihood (Fig 5, blue). Removing any one of its components results in a significant decrease in quality of fit (Fig 5, red), while adding additional components of any of the three types does not significantly increase quality of fit (Fig 5, green).

**Table 1:** Parameter equivalencies between a mixture-of-exponentials model with three components and the cognitive model.

| Parameter Equivalencies |  |
| --- | --- |
| Mixture of Exponentials Model | Cognitive Model |
| $\alpha_{v,1}; \alpha_{h,1}$ | $\alpha_v; \alpha_h$ |
| $\alpha_{v,2}; \alpha_{h,2}$ | $\alpha_g$ |
| $\beta_{v,1}; \beta_{h,1}$ | $\beta_v; \beta_h$ |
| $\beta_{v,2}; \beta_{h,2}$ | $\beta_g$ |
| $\beta_{bias}$ | $\beta_{bias}$ |

**Figure 5:**
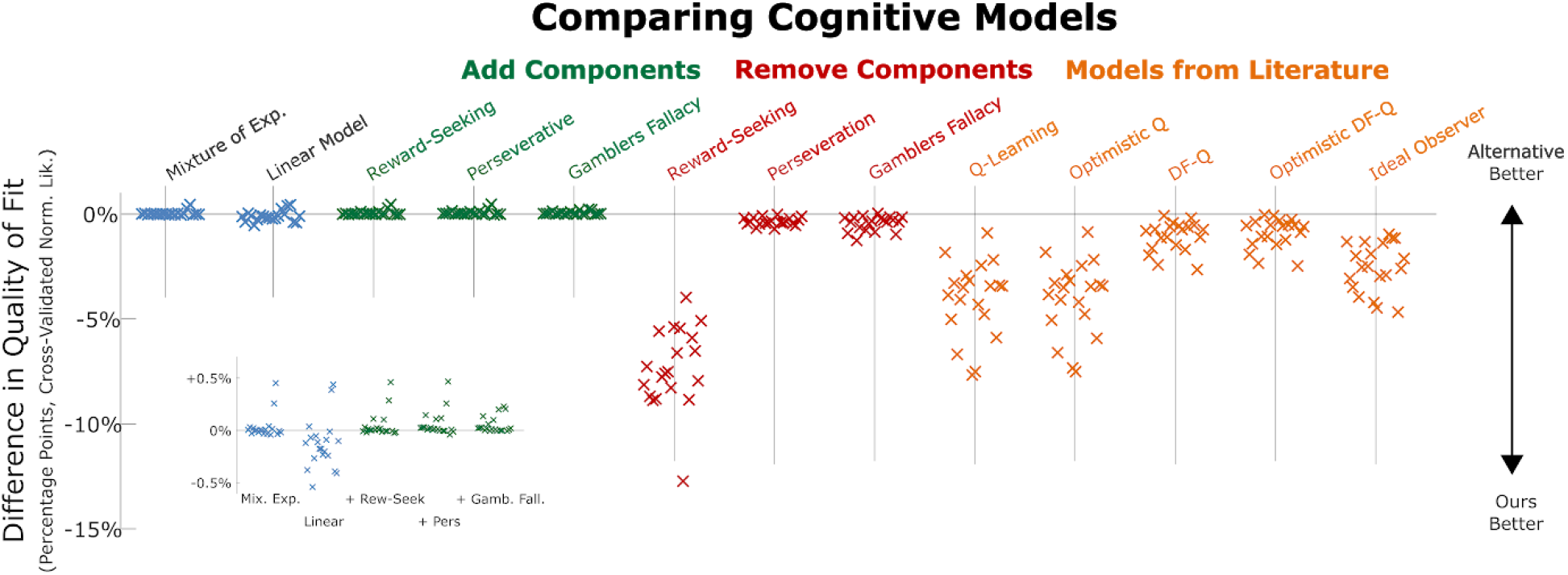
Cognitive Model Comparison. Difference in normalized cross-validated likelihood between our cognitive model and various alternatives. The first two alternatives (blue) are models described earlier in the paper: the linear model and the mixture of exponentials model. The next three alternatives (green) are versions of our model that have been expanded to include an extra copy of one of the three components. The next three alternatives (red) are versions of our model that have been reduced by removing one of the three components. Finally are five models from the literature (orange): Q-Learning (e.g. Kim et al., 2009), Optimistic RL (Lefebvre et al., 2017), Differential Forgetting Q-Learning (Ito & Doya, 2009), a hybrid combining both optimistic and differential forgetting mechanisms, and a Bayesian ideal observer model (Daw et al., 2006). The inset shows an expanded view of the first five models.

### Comparison to Existing Models

Reward learning tasks like ours have been extensively studied in behavioral neuroscience, and many cognitive models have been used to interpret behavioral and neural data. We evaluated several of these to see whether they could provide a better fit than our cognitive model (Figure 5, orange). The first was a q-learning model, variants of which have been used in a wide variety of tasks and species (Bari et al., 2019; e.g. Barraclough et al., 2004; Kim et al., 2009). This model is similar to the reward-seeking component of our model in that it maintains internal variables related to reward history, but different in that it uses separate variables to track reward history at each port, which are updated only on trials where their port is visited. The second (“optimistic RL” Lefebvre et al., 2017) is an extension to the q-learning model in which different learning rates are applied to better-than-expected and to worse-than expected outcomes. The third (“differential forgetting” Ito & Doya, 2009) is an alternative extension, which updates estimates both for visited and unvisited ports, using different learning rates, and also applies different learning targets in the case of positive and negative outcomes. Fourth was a hybrid model containing both optimistic and differential forgetting extensions. Finally, we also considered an ideal observer model, which uses a Bayesian filter to update a belief distribution over the current reward probabilities (Daw et al., 2006), and selects actions based both on immediate expected reward and on reduction of uncertainty (Wilson et al., 2014). We evaluated each of these models using two-fold cross validation, and found that our three-component cognitive model provided a better fit than any of them, for all rats in our dataset (n=20; p=10^-4^, signrank test). This validates the idea that our approach of successive model reduction can result in cognitive models that compare favorably, in terms of predictive performance, with others in the literature.

## Discussion

We trained rats to perform a reward-learning task in which they repeatedly selected between two ports with constantly-changing probabilities of reinforcement, and we sought to build a cognitive model capturing the patterns in their behavior. This work adds to a large body of previous work modeling behavior in tasks of this kind (Daw et al., 2006; Ito & Doya, 2009; Kim et al., 2009; Samejima et al., 2005), building on it by collecting a large behavioral dataset, and by adopting a data-first modeling approach to reveal behavioral patterns in a hypothesis-neutral way.

In keeping with this literature, we built models which sought to predict the choice of the subject on each trial, using as predictors the history of choices made and rewards received on recent trials. In making this decision, we choose to ignore (to treat as noise) all other factors that might influence choice, including factors such as satiety or fatigue that may play an important role in modulating behavior. We began by fitting the most flexible models possible that are consistent with this choice, the unconstrained models. Because of their flexibility, these models provide a ceiling on the quality of fit achievable by any model that considers the same inputs as they do. In the regime where the unconstrained model does not overfit, this ceiling is also a useful benchmark – any model which accurately captures the generative process should fit the data approximately as well as the unconstrained model does. An important precedent to this benchmarking step is found in Ito & Doya (2009), who similarly fit unconstrained models, and compared them to several cognitive models. Our work adds to this by comparing training-dataset and testing-dataset performance to determine the range in which the benchmark established by these models is valid, and by utilizing a larger dataset, which allows us to establish a benchmark for models considering longer histories.

With this benchmark in hand, we fit a set of models with an additional constraint of linearity – these models assume that the outcome of each past trial contributes independently to present choice. The performance of the linear models matches the benchmark established by the unconstrained models, suggesting that their linearity assumption is a valid one. These linear models have an important precedent in earlier work (Corrado et al., 2005; Lau & Glimcher, 2005), which used similar models to characterize behavioral data from primates performing reward-guided decision tasks. Our work builds on this in that it validates these models with respect to the benchmark from the unconstrained models, and constructs models that are further reduced, including one that can be viewed as a cognitive model.

Fits of the linear model indicate that rats have strong tendencies of “reward-seeking” (repeat choices that lead to reward, switch from those that do not) and “perseveration” (repeat past choices without regard to outcome). We sought to compress these patterns using a model that assumes they arise from a mixture of several exponential processes with distinct weights. We found that the dataset from each rat was best modeled as arising from a mixture of exactly two exponential processes of each type, and that the parameters of these showed consistent patterns across rats. Specifically, the slower reward-seeking and the slower perseverative process had learning rates that were nearly equal, and had weights that were nearly equal-and-opposite. Finally, we wrote a cognitive model that captured these patterns using a mixture of agents model. This type of model has important precedents in mixture-of-agents models for related tasks, including working-memory/reinforcement-learning (Collins & Frank, 2012), episodic-memory/reinforcement-learning (Bornstein & Daw, 2012) and model-based/model-free (Daw et al., 2011). Our cognitive model comprised a mixture of three agents. The fastest was a reward-seeking agent, while the middle process was a perseverative “habits” agent, and the slowest was a “lose-stay” agent that tends to repeat choices that have led in the past to losses.

The first, reward-seeking, component of the model is consistent with a long body of literature on reward-guided decision making, beginning with Thorndike’s “law of effect”, which holds that actions that have led to reward in the past are likely to be repeated in the future (Thorndike, 1911). It is reminiscent of popular reinforcement learning algorithms, such as the delta rule (Sutton & Barto 2017, which are commonly used to model behavior on tasks of this kind (Bari et al., 2019; Barraclough et al., 2004; Daw et al., 2006; Kim et al., 2009). These algorithms are similar to our exponential process in that they maintain a cached variable, updating it on each trial by taking its weighted average with a value from that trial’s observation. They are different in that they typically maintain one cached value per possible action, while our process caches only a single value, representing the relative history of recent outcomes at each port.

Perseveration in reward-guided tasks has been observed in a wide variety of tasks and species (Balcarras et al., 2016; Ito & Doya, 2009; Kim et al., 2009; Lau & Glimcher, 2005; Lee et al., 2005; Rutledge et al., 2009). One common way of modeling it is to consider it as arising from some modulation to a reward-seeking process – for example using different learning rates for positive and negative outcomes (Ito & Doya, 2009; Lefebvre et al., 2017), or by fitting a perseverative weight that considers only the immediately previous trial (Daw et al., 2011). Our results suggest that perseveration results from a process that is separable from reward-seeking (having its own timecourse), and that it considers a relatively large number of past trials. The idea of a separable perseverative process is consistent with Thorndike’s second law, the “law of exercise”, which holds that actions which have often been taken in the past are likely to be repeated in the future (Thorndike, 1911). It is also consistent with ideas from the psychology of habit formation (Dickinson, 1985; Graybiel, 2008; Wood & Rünger, 2016), which propose that habits are fixed patterns of responding that result from simple repetition of past actions (Ashby et al., 2010; Miller et al., 2018, 2019). While we are not aware of other models for bandit tasks that incorporate this mechanism, a very similar one has been proposed to play a role in sensory decision-making (Akaishi et al., 2014).

The “lose-stay” pattern of repeating choices that have led in the past to losses is reminiscent of the “gambler’s fallacy”, in which humans seem to believe that a series of losing outcomes predicts that a winning outcome is likely imminent. This pattern occurs when humans attempt to predict the next outcome of a process that they believe to be random, such as the flip of a coin or the spin of a roulette wheel (Oskarsson et al., 2009). While no clear consensus exists as to the cause of the gambler’s fallacy, several explanations have been proposed. Perhaps the oldest proposes that humans believe in a “law of small numbers”, expecting that even very short sequences will include an equal number of winning and losing outcomes, and that a surplus of losses in the past must therefore be balanced out by an increased proportion of wins in the future (Tversky & Kahneman, 1971). This is proposed to be an example of the more-general “representativeness heuristic”, in which people expect each instance from a category to embody the properties of that category as a whole. More recent accounts propose that humans have complex, if often erroneous, internal models of what to expect from “random” processes, and that these give rise to the gambler’s fallacy (Oskarsson et al., 2009). A final proposal is that the gambler’s fallacy results from a rational bias that arises when a resource-limited learning system attempts to learn from the true statistics of random sequences (Hahn & Warren, 2009; Sun et al., 2015). None of these proposals seems to us to provide an immediate explanation of our lose-stay agent, but they do raise hope that some related cognitive explanation might be possible. To our knowledge, previous studies using similar tasks have not reported a pattern of this kind. It is possible that it is specific in some way to our experimental setup (e.g. specific to rats, specific to nose-ports, specific to fluid rewards, etc). It is also possible that it is present broadly in behavior on tasks of this kind, but has not previously been identified on account of being unexpected and somewhat subtle.

This three-agents cognitive model has a number of features that make it a reasonable hypothesis about the algorithm actually used by the brain to perform our task. Each of its agents (reward-seeking, perseveration, and lose-stay) has a relatively simple implementation, with just two free parameters that govern the evolution of just one hidden variable, and is cognitively plausible given the rest of what is known about the psychology and biology of behavior on tasks of this kind. It makes several claims that are distinct from those of other cognitive models in the field. One of these is that the reward-seeking component of behavior contains a single latent variable that is influenced by rewards following both choices, rather than separate latent variables for each available action. Another is that there is a separable lose-stay (gambler’s fallacy) component of behavior. This model also provides a better quantitative fit to our dataset than do other cognitive models drawn from the literature. We produced this model by a process of successive model reduction: beginning with a highly general model that provides a good fit to data but is not itself cognitively plausible, identifying patterns in the fit parameters of this model, writing a simpler model that embodies those patterns as structural assumptions, and repeating. We believe that it validates the idea that this approach of successive reduction is capable of producing useful cognitive models.

## Acknowledgements

We would like to thank Stefano Palminteri, Alex Piet, Kim Stachenfeld, Nathaniel Daw, Yael Niv, Bas van Ophuesden, Chuck Kopec, and Jeff Erlich for helpful discussions, and Kim Stachenfeld, Alex Piet, and Anna Lebedeva for helpful comments on the manuscript.

## Methods

### Subjects

All subjects were adult male Long-Evans rats (Taconic Biosciences), placed on a restricted water schedule to motivate them to work for water rewards. Rats were housed on a reverse 12-hour light cycle and trained during the dark phase of the cycle. Rats were pair housed during behavioral training. Animal use procedures were approved by the Princeton University Institutional Animal Care and Use Committee and carried out in accordance with NIH standards.

### Behavioral Apparatus

Rats were trained in custom behavioral chambers (Island Motion, NY) located inside sound- and light-attenuated boxes (Coulbourn Instruments, PA). Each chamber was outfitted with three nose ports arranged horizontally. Each port contained a white light emitting diode (LED) for delivering visual stimuli, as well as an infrared LED and phototransistor for detecting rats’ entries into the port. The left and right ports also contained sipper tubes for delivering water rewards. Training was controlled by a custom protocol written using the bControl behavioral control system. Rats were placed into and removed from training chambers by technicians blind to the experiment being run.

### Two-Armed Bandit Task: Training Pipeline

Here we outline a procedure suitable for efficiently training naive rats on the two-armed bandit task. Automated code for training rats using this pipeline via the bControl behavioral control system will be made available on the Brody lab website (http://brodylab.org/). This formalization of our training procedure into a software pipeline should also facilitate efforts to replicate our task in other labs, as the pipeline can readily be downloaded and identically re-run.

#### Phase I: Sipper tube familiarization

Training began with an initial period of sipper tube familiarization. In each trial of this phase, the LED in either the left or the right port would illuminate, and a 50 uL reward would be delivered if the rat entered this port. Training in this phase continued until the rat was completing approximately 200 trials per day. Typically, training in this phase requires three to five sessions.

#### Phase II: Trial structure familiarization

Training in this phase typically lasted three to seven days. In the second phase, each trial began with illumination of the central port’s LED. When the rat entered this port, its LED would extinguish, and the LED in either the left or the right port would illuminate. Entry into this port resulted in reward. Training continued until the rat was completing approximately 200 trials per day.

#### Phase IIIa: Performance-triggered flips

In this phase, decision-making and dynamic reward contingencies are introduced. Trials began with illumination of the central port’s LED. In 90% of trials (“free choice trials”), both the left and right port’s LEDs would illuminate and the rat could enter either. In half of the remaining trials (5% of total), only the left port’s LED would illuminate, and in the other half, only the right port’s LED would illuminate. In these “instructed choice” trials, the rat was required to enter the lit port in order for the task to continue. In all trial types, reward contingencies were dynamic and changed in blocks. In each block, either the left or the right port was “good”, while the other was “bad”. Entry into the good port resulted in reward, while entry into the bad port resulted only in the initiation of a new trial. Whether the left or the right port was good was initialized randomly at the beginning of each session, and reversed if the rat selected the good port on more than 80% of the previous 50 free choice trials. Training in this phase continues until the rat achieves a large number of flips per session. Typically this requires only one or two sessions.

#### Phases IIIb and IIIc

These phases are the same as phase IIIa, except that the good and bad ports are rewarded 90% and 10% of the time in phase IIIb, and 80% and 20% of the time in phase IIIc. Training in each phase continues until the rat achieves a large number of block changes per session. Typically this requires two to five sessions.

#### Final Task: Drifting Two-Armed Bandit Task

In the final task, reward probability of the two ports no longer depends on the behavior of the rat, and no longer shifts in blocks. Reward probability of the left and right port are sampled randomly between zero and one at the start of each session, and are updated after each trial following a Gaussian random walk. Specifically, the reward probability is sampled according to P_t+1_ ~ Gaussian(μ=P_t_, σ=0.15), where *P_t_* is the probability of reward on trial *t*. If this would result in a probability greater than one (less than zero), *P_t_* is set to one (zero) instead.

### Model Fitting and Comparison

All model fitting was done using custom code written in Matlab (Mathworks). In all cases, models were fit by maximum-likelihood or maximum-a-posteriori, with separate fits for each rat. Unconstrained models (n-markov models) were fit according to equations one and two (see Main Text). Linear models were fit using the Matlab function `glmfit`, implementing unregularized logistic regression. Mixture-of-exponentials models and cognitive models were fit using custom likelihood functions written in the probabilistic programming language Stan (B. Carpenter et al., 2016), accessed using the MatlabStan (Stan Development Team, 2016) interface. Mixture-of-exponentials models were fit using weakly informative priors: a `beta(3,3)` prior for learning rate parameters, and a `gaussian(0,1)` prior for weighting parameters.

For analysis reporting fit parameters (Fig. 2ab, Fig. 3a, Fig. 4bc, Supplemental Figures), models were fit to the entire dataset for each rat. For analysis reporting model comparison (Fig. 2c, Fig. 3bc, Fig. 4a, Fig. 5) each model was fit twice – once to behavioral data from even-numbered sessions, and once to odd-numbered sessions. The likelihood of each model was evaluated both on the dataset that it was fit to (even or odd; “training dataset”), and once to the other dataset (odd or even; “testing dataset”). To compare likelihood across rats, we compute “normalized likelihood” (Daw, 2011)

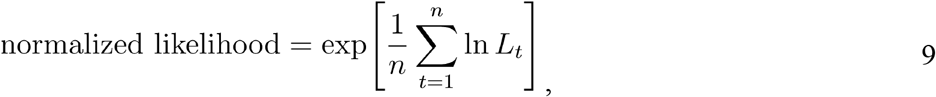

where *L_t_* indicates the likelihood the model assigned to the choice that the rat actually made on trial *t*, *n* indicates the total number of trials performed by the rat. This can be interpreted as the geometric mean over the probabilities with which the model would have taken the actions which the rat actually took.

### Q-Learning Models

We compared our cognitive model (equations 5–8) to several others from the literature (Fig 4). Many of these are variants of q-learning models, and can be written as special cases of a “hybrid” model, each with different parameter constraints. The hybrid model maintains separate estimated values for the left and for the right port (*Q_left_* and *Q_right_*), both of which are initialized to zero at the beginning of each session. One each trial, the model selects a port according to a softmax decision rule based on Q:

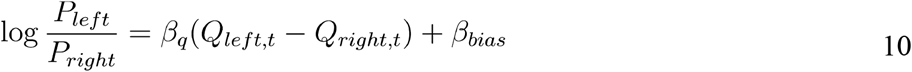

where *β_Q_* and *β_bias_* are weighting parameters (also called “inverse temperature” parameters) governing the relative importance of estimated port values, which vary trial by trial, and of a fixed left/right bias. After each trial, the estimated value of the chosen and unchosen ports are updated according to:

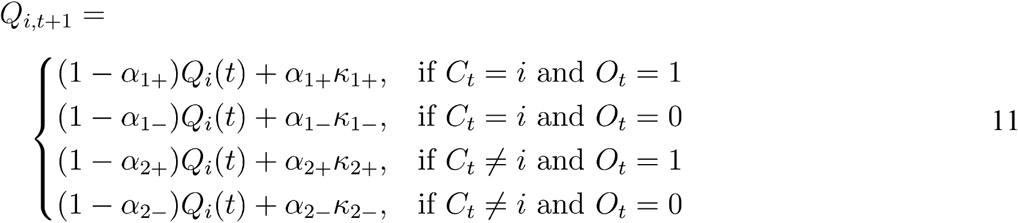

Where *C_t_* is the action taken on trial *t* (left or right), *O_t_* is the outcome received on trial *t* (reward or omission), the *α* parameters represent learning rates, and the *κ* parameters represent learning targets. Parameters with a subscript 1 govern update of the estimated value of the port that was chosen, while those with subscript 2 govern update of the estimated value of the unchosen port. Parameters with subscript + govern updates following rewarded trials, while those with subscript - govern updates following unrewarded trials.

Of the models we consider using this notation, the “Q learning” model is the most constrained. This model does not update the estimated value of the unchosen port (α_2+_ = α_2-_ = *κ*_2+_ = *κ*_2-_ = 0). It also uses the same learning rate for rewarded and for unrewarded trials (α_1+_ = α_1-_), and constrains the learning targets (*κ*_1+_= 1; *κ*_1-_= 0). This model has three free parameters: *β_Q_*, *β_bias_*, and *α*_1_.

“Optimistic RL” (Lefebvre et al., 2017) relaxes some of these constraints, allowing different learning rates on rewarded and on unrewarded trials, allowing *α*_1+_ and *α*_1-_ to separately vary. It retains the constraints on learning about the unchosen port (*α*_2+_ = *α*_2-_ = *κ*_2+_ = *κ*_2-_ = 0), and on the learning targets (*κ*_1+_= 1; *κ*_1-_= 0). This model has four free parameters: *β_Q_*, *β_bias_*, *α*_1+_, and *α*_1-_.

“Differential Forgetting Q-learning” (Ito & Doya, 2009) relaxes a different set of constraints. This model allows learning targets for the chosen option (*κ*_1+_ and *κ*_1-_) to vary freely. It also allows “forgetting”, in which the value of the unchosen option is updated. This involves allowing *α*_2+_ and *α*_2-_ to take values other than zero. In this model, forgetting is identical between rewarded and unrewarded trials (*α*_2+_ = *α*_2-_), and always has a target of 0 (*κ*_2+_ = *κ*_2-_ = 0). The DF-Q model also constrains *β_Q_*=1, since this parameter is redundant with *κ*_1+_. This model has five free parameters: *β_bias_*, *α*_1_, *α*_2_, *κ*_1+_ and *κ*_1-_.

Finally, we also consider the complete hybrid model given by equations 10 and 11. This model has ten free parameters: *β_Q_*, *β_bias_*, *α*_1+_, *α*_1-_, *α*_2+_, *α*_2-_, *κ*_1+_, *κ*_1-_, *κ*_2+_, and *κ*_2-_ Some of these parameters are redundant with one another.

All q-learning models were instantiated and fit using the probabilistic programming language Stan (Bob Carpenter et al., 2017) via its Matlab interface (Stan Development Team, 2016). Models were fit by maximum-a-posteriori, using weakly informative priors of `beta (3, 3)` prior for learning rate parameters (*α*), and `gaussian (0, 1)` for weighting and target parameters (*β* and *α*).

### Ideal Observer Models

We also compared our cognitive model to an ideal observer model. This model implements a Bayesian filter, which maintains a belief distribution over the reward probabilities of each port. We use *θ_left_* and *θ_right_* to parameterize the reward probability on the left and on the right port, and *B_t, right_* and *B_t_* indicate the observer’s belief at time *t* over these probabilities. We approximate these distributions numerically, allowing theta to take 101 discrete values equally spaced between zero and one. Belief distributions are initialized to be uniform at the start of each session, and updated twice following each trial. The first is a Bayesian update applied to the belief distribution for the chosen port:

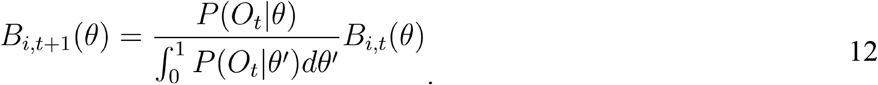

where *B_i_* is the belief distribution associated with the port chosen on trial *t*, *O_t_* is the outcome observed on trial *t*, and *θ* parameterizes the probability of reward at that port. Because of this, *P*(*O_t_*|*Θ*) = *Θ* on rewarded trials, and *P*(*O*_t_|*Θ*) = (1-*Θ*) on unrewarded trials. The second update accounts the trial-by-trial drift in reward probabilities. Each belief distribution is convolved with a gaussian kernel whose standard deviation (*σ*) is a free parameter corresponding to the observer’s belief about the environmental drift rate.

To make choices, the ideal observer first computes the mean (*V*) and variance (*U*) of the reward probabilities given its current belief distributions, which quantify the expected immediate reward given its current beliefs, as well as the uncertainty in that expectation:

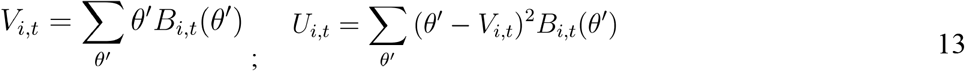

The decision on each trial was made using a weighted average of the differences in these quantities:

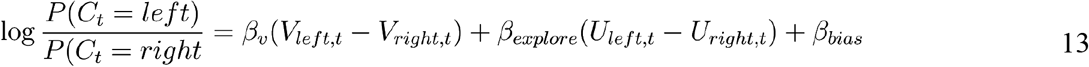

where *β_v_* quantifies the model’s tendency to select the port with the currently higher expected reward probability, *β_explore_* quantifies a tendency to choose the side with currently greater uncertainty in reward probability ("directed exploration"; Daw et al., 2006; Wilson et al., 2014), and *β_bias_* quantifies a fixed bias towards the left or the right port. The model therefore had four free parameters: these four *β*s, as well as *σ*. For quantitative model comparison (Figure 4), we computed normalized likelihoods using two-fold cross-validation, as described above, and maximum likelihood fits using code written in Matlab. For generating synthetic data (Figure 1c), we set σ to its true generative value of 0.15, β_*v*_ to 100, and both *β_explore_* and *β_bias_* to zero.

### Software and Data Availability

Automated code for training rats on the two-armed bandit task will be made available on the Brody lab website upon publication. Software used for data analysis, as well as processed data will be made publicly available upon publication. Raw data are available from the authors upon request.

---

**Figure S1:**
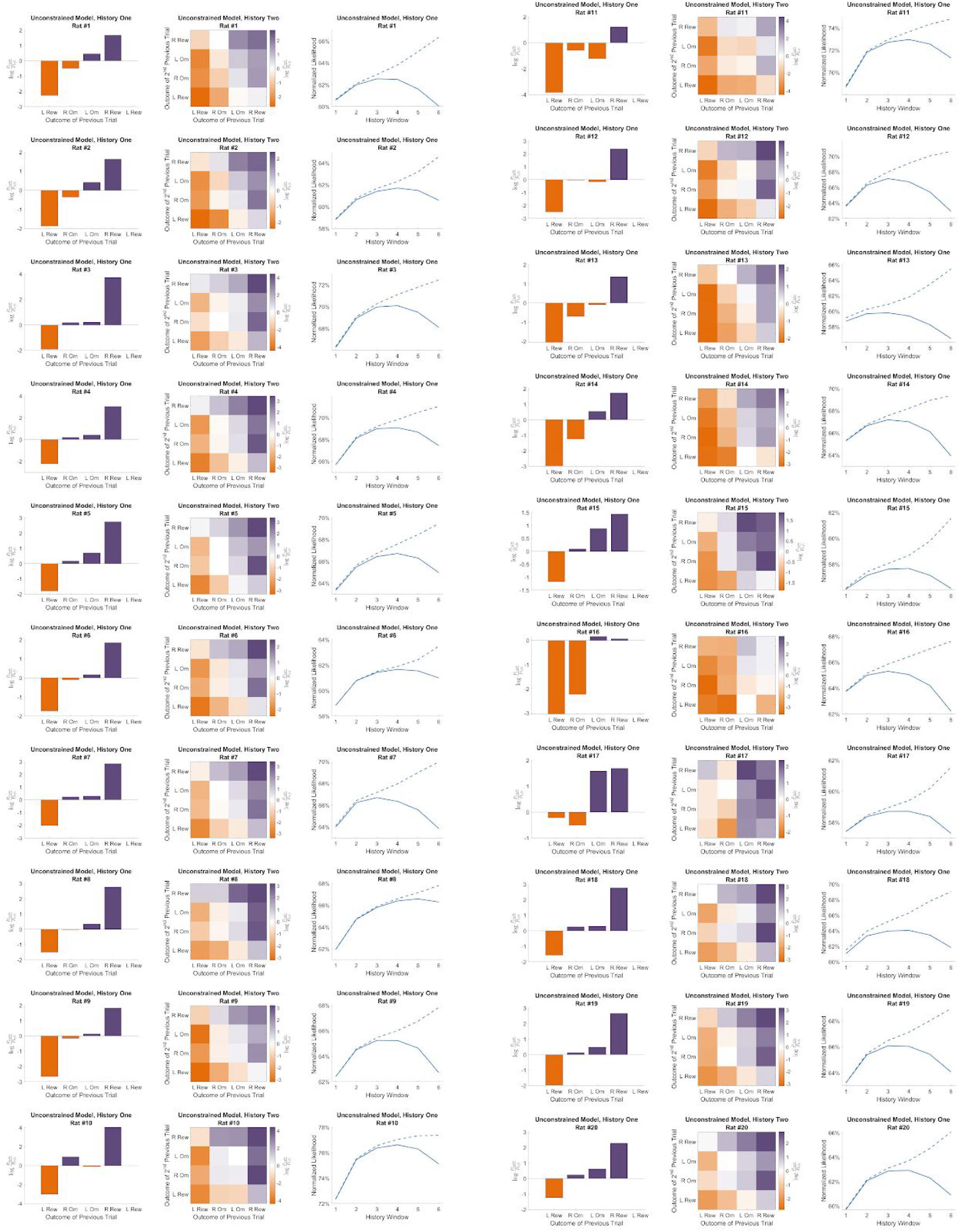
Fits of the unconstrained model for all rats.

**Figure S2:**
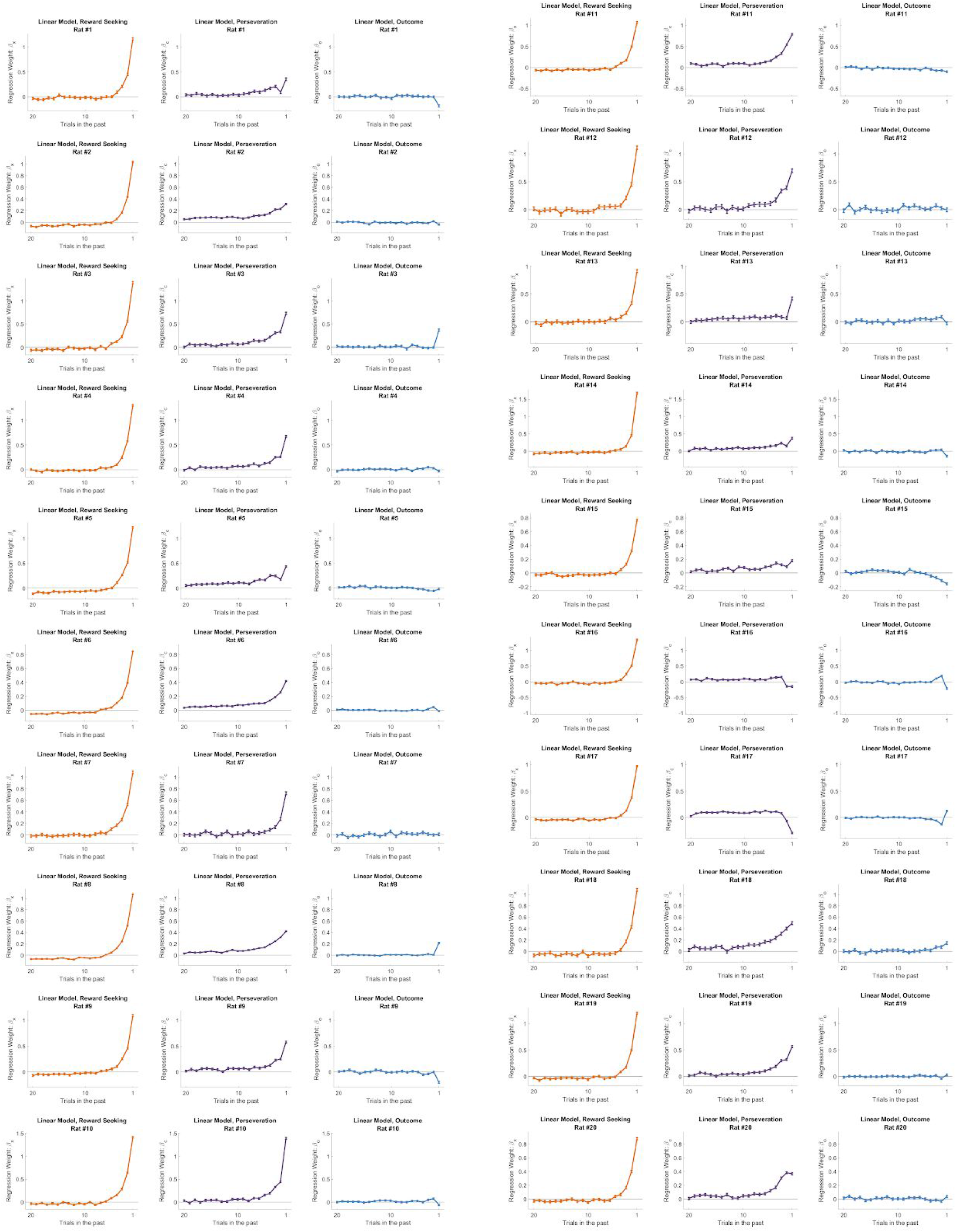
Fits of the linear model for all rats.

